# Disentangling the contribution of childhood and adulthood circumstances and genetics to phenotypic aging: prospective cohort study

**DOI:** 10.1101/384040

**Authors:** Zuyun Liu, Xi Chen, Thomas M. Gill, Chao Ma, Eileen M. Crimmins, Morgan E. Levine

## Abstract

**Objectives:** To evaluate the extent to which childhood and adulthood circumstances and genetics contribute to phenotypic aging, using a multi-system-based signature of aging that has been shown to capture mortality and morbidity risk.

**Design:** Prospective population-based cohort study.

**Setting:** United States (U.S.).

**Participants:** 2,339 adults (aged 51+ years) from U.S. Health and Retirement Study, who participated in the Core Survey, the 2016 Venous Blood Study, the 2015 Life History Mail Survey, the Enhanced Face-To-Face interview (2006-2016), and were part of the genetic sample.

**Main outcomes measure:** Phenotypic Age, a validated aging measure based on a linear combination of chronological age and nine multi-system biomarkers. For most analyses, we examined “PhenoAgeAccel”, which represents phenotypic aging after accounting for chronological age (i.e. whether a person appears older [positive value] or younger [negative value] than expected, physiologically).

**Results:** The Shapley Value Decomposition approach revealed that together all 11 domains (four childhood and adulthood circumstances domains, five polygenic scores [PGSs] domains, demographics, and behaviors domains) accounted for about 30% of variance in PhenoAgeAccel. Among the four circumstances domains, adulthood adversity was the largest contributor (9%), while adulthood socioeconomic status (SES), childhood adversity, and childhood SES accounted for 2.8%, 2.1%, 0.7%, respectively. Collectively, all PGSs contributed 3.8% of variance in PhenoAgeAccel. Further, six subpopulations/clusters—identified using a hierarchical cluster analysis based on childhood and adulthood SES and adversity—showed differences in average levels of phenotypic aging. Finally, there was a significant gene-by-environment interaction between a previously validated PGS for coronary artery disease and the most apparently disadvantaged subpopulation/cluster—suggesting a multiplicative effect of adverse environment coupled with genetic risk on phenotypic aging.

**Conclusions:** Socioenvironmental circumstances during both childhood and adulthood account for a sizable proportion of the difference in phenotypic aging among U.S. older adults. The detrimental effects may further be exacerbated among persons with a genetic predisposition to coronary artery disease.

## INTRODUCTION

Aging is a complex multifactorial process, characterized by increasing dysregulation and loss of function across multiple levels and systems.^1^ Consequently, the aging process is presumed to be a major driver in the pathogenesis of many chronic diseases.^2 3^ While the process of aging is universal, individuals are heterogeneous in their rates of aging, which in turn, directly influence susceptibility to morbidity and mortality events—faster aging is reflected in earlier incidence of disease and death.^4–6^ Thus, developing behavioral, social, or pharmacological interventions that slow the aging process will directly impact population health.

Aging begins at conception; thus, interventions will be most successful if applied early in the life course, prior to the onset of disease and disability.^7^ However, to be feasible, one needs to employ “biomarkers of aging”, which can both identify at-risk individuals who show signs of accelerated aging, and evaluate intervention efficacy. Phenotypic Age (PhenoAge) is a multi-system-based signature of aging that we have developed and validated in large nationally representative U.S. samples.^8^ ^9^ This signature has been shown to better predict all-cause and disease-specific mortality than chronological age, even among healthy individuals. PhenoAge is meant to capture age-related dysregulation and predict subsequent health declines. While we know that differences in this signature translate into variations in morbidity/mortality susceptibility, we have yet to disentangle the contribution of potential factors to differential phenotypic aging.

Previous work has provided strong evidence that traumas and adversities in childhood and adulthood influence risk of various outcomes (e.g. disease and mortality) in later life,^10–13^ presumably via an acceleration of the aging process.^14–17^ The cumulative “wear and tear” in response to chronic stressors or deprivation experienced over the life course is thought to “get under the skin” by contributing to declines in physiological adaptation (i.e. allostatic load) that manifest as vulnerability to disease and death.^18 19^ Socioeconomic status (SES) is also thought to be a powerful driver of health and aging disparities.^14 16 20–28^ Gaps in wealth and education between the “haves” and “have nots”—particularly in the U.S.—have been shown to produce stark differences in risk for long-term aging-related outcomes.^14 16 28^ To capture a more complete picture of how various factors coalesce and in turn manifest as health disparities, it is necessary to assemble a comprehensive set of circumstances that can be examined concurrently.

In addition to socioenvironmental circumstances, genetics contribute to differential vulnerability for aging and disease. For example, it has been estimated that about 20-30% of lifespan is genetically determined.^29^ Similarly, twin studies have estimated that fatal coronary heart disease is about 40-50% heritable,^30–32^ while many cancers have heritability estimates around 30-60%.^33^ Therefore, accounting for innate differences in genetic susceptibility becomes important when estimating the influence of the environment for health and aging traits.^34^ Moreover, genetics may also have the potential to alter an individual’s vulnerability to various environmental conditions. For instance, a gene-by-environment interaction (GxE) is defined as “a different effect of an environmental exposure on disease risk in persons with different genotypes.”^35^ Under this assumption, adverse socioenvironmental exposures—such as low SES and adversity—may be more detrimental to a person’s health if s/he is already genetically at-risk of death and/or disease outcomes.

While various factors, including socioenvironmental circumstances and genetics, influence health and aging to some extent, their relative contributions to aging and disease risk are uncertain. The multitude of factors defining an individual’s specific life circumstances poses challenges for modeling their individual and cumulative effects. Another challenge is how to differentiate the effects of circumstances in childhood from those in adulthood. From the perspective of health economics, variance in health (i.e. inequality) can be influenced by circumstances throughout the life course; yet, there are variations in the level of control (modifiability) a person has over the circumstances shown to contribute to health and aging inequalities. For instance, one’s demographics, genetics, and childhood SES and experiences of traumas are essentially a lottery, yet are all believed to have down-stream consequences for health and aging. This has been referred to as Inequality of Opportunity.^36 37^ Furthermore, it is important to note that an individual’s circumstances in adulthood can be strongly influenced/confounded by his/her circumstances as a child, or even genetic predisposition for various personality and mental health traits and/or educational attainment.^38^ One key contribution of this research angle is to innovatively discern the effect of circumstances beyond one’s control, which should be the priority of public policies that aim to alleviate health inequality. The Shapley Value Decomposition approach facilitates estimation of the share of health inequality due to circumstances with varying degrees of modifiability. By appropriately assigning contributions from sources of health inequality, the Shapley Value Decomposition approach estimates the overall and relative importance with substantial advantages (see more in supplementary materials appendix 1).

The present study assembled a comprehensive set of variables assessing childhood and adulthood circumstances and genetics using a nationally representative sample of older adults in the U.S. By innovatively applying two approaches—Shapley Value Decomposition and Hierarchical Clustering—we were able to disentangle the relative and cumulative contributions of these various factors to variations in aging and morbidity/mortality risk, as captured using our novel PhenoAge measure.

## METHODS

### Data

The Health and Retirement Study (HRS) is an ongoing nationally representative, biennial survey of older Americans (aged 51+ years), and their spouses, beginning in 1992.^39^ HRS is funded by the National Institute on Aging and carried out by the University of Michigan. In this study, we assembled a large array of variables from four data sources (sub-studies) within HRS, including the core survey (1996-2016), the newly released 2015 Life History Mail Survey (LHMS), the Enhanced Face-To-Face (EFTF) interview (2006-2016), and the 2016 Venous Blood Study (VBS). A description of the four data sources can be found in supplementary materials and elsewhere.^39^ After restricting to persons who participated in each of these sub-studies, our final analytic sample included 2,339 persons (Figure S1). Compared with persons who participated in both the 2015 LHMS and 2016 VBS but were excluded in this analysis, our analytic sample showed a similar sex ratio but were older (69.4 vs 68.3 years), more highly educated (13.9 vs 13.2 year), and more likely to be non-Hispanic white (93.9% vs 84.0%).

### Participant involvement

According to the nature of the dataset and the ethical permission, participants were not involved in setting the research agenda, nor were they involved in developing plans for the design or implementation of this study. There are no plans to directly disseminate the findings of this study to participants, but dissemination to the general public and peers will be undertaken by using presentations and social media.

### Childhood and adulthood circumstances

Since no consensus has been reached regarding the selection and definitions of circumstance variables, we considered a comprehensive set of measures based on literature suggesting their potential relationship with health- and/or age-related outcomes. All questions and corresponding responses/descriptions are provided in Table S1. In brief, we defined four domains of childhood and adulthood circumstances: childhood SES, childhood adversity, adulthood SES, and adulthood adversity.

### Genetic factors

To further differentiate the effect of circumstances on phenotypic aging due to genetic predisposition (i.e., beyond one’s control), we included five domains for genetic factors based on previously established polygenic scores (PGSs): anthropometrics, disease/longevity, mental health/personality, education/cognition, and smoking (details can be found in supplementary materials appendix 1). The saliva samples for genotyping SNPs were collected in the EFTF interview from 2006 to 2012, and details on the construction of these PGSs are provided elsewhere.^40^

### PhenoAge and phenotypic aging (PhenoAgeAccel)

PhenoAge was first developed and validated using independent waves from the National Health and Nutrition Examination Survey (NHANES).^9^ In brief, PhenoAge was derived from nine biomarker variables, selected out of a possible 42 using an elastic net proportional hazards model for mortality. These biomarkers included albumin, creatinine, glucose, (log) C-reactive protein, lymphocyte percent, mean cell volume, red cell distribution width, alkaline phosphatase, and white blood cell count. Chronological age was also included. The score was calculated as a weighted (coefficients) linear combination of these variables, that was then transformed into units of years using two parametric (Gompertz distribution) proportional hazard models—one for the linearly combined score for all ten variables and another for chronological age. Thus, PhenoAge represents the expected age within the population that corresponds to a person’s estimated hazard of mortality as a function of his/her biological profile.^8 9^

Next, we calculated a measure of “age acceleration” (i.e., PhenoAgeAccel), defined as the residual resulting from a linear model when regressing PhenoAgeAccel on chronological age. Therefore, PhenoAgeAccel represents phenotypic aging after accounting for chronological age (i.e. whether a person appears older [positive value] or younger [negative value] than expected, physiologically).

### Statistical Analyses

More complete details of the analytic plan can be found in the supplementary materials appendix 1. Briefly, we first used the Shapley Value Decomposition approach with mean logarithmic deviation (MLD) method to evaluate the overall and relative contributions of all variables including childhood and adulthood circumstances and genetics to PhenoAgeAccel. Compared with other decomposition methods, the Shapley Value Decomposition approach has substantial advantages, such as being order independent (i.e. the order of circumstances for decomposition does not influence the results) and being able to sum components to produce the total value.

To potentially inform possible intervention strategies, we then assessed the associations of childhood and adulthood circumstances with PhenoAge or PhenoAgeAccel using a series of analyses including clustering analysis. First, we performed a principal component analysis for the four domains (childhood SES, childhood adversity, adulthood SES, and adulthood adversity) that are potentially correlated. Second, based on the first four principal components, selected via a series of assessments, we performed a hierarchical clustering analysis (HCA) to categorize participants into distinct subpopulations/clusters, representing groups of people with shared life experiences. To make it clearer, we calculated a continuous measure (i.e. cluster membership, ranging from −1 to 1) for each cluster that denotes how similar a participant’s profile is to the characteristics represented by the cluster. This was estimated as the correlation between the participant’s scores across the variables used for clustering, and the first principal component when only considering persons assigned to the cluster. For instance, someone may have a score of 0.8 for cluster 1 and −0.6 for cluster 2, suggesting s/he is very similar to the profile representative of cluster 1, but not cluster 2. We then related these cluster membership scores to the circumstances measures to determine what characteristics define each cluster. Third, we compared the PhenoAgeAccel of participants assigned to each subpopulation/cluster. Fourth, we examined the association of these subpopulations/clusters with PhenoAge using ordinary least squares (OLS) models with adjustment for covariates such as chronological age, sex and ancestry. Finally, we examined GxE interactions, by testing whether the PGS (selecting the most significant one) further increased differences in PhenoAge between the subpopulations/clusters.

## RESULTS

Sample characteristics are shown in Table S2 and described in supplementary materials.

### The contribution of childhood and adulthood circumstances and genetics to phenotypic aging

Figure 1 presents the results from the Shapley Value Decomposition approach, depicting the proportions of the variance explained by all 11 domains. Collectively, the factors evaluated accounted for about 30% of the variance in PhenoAgeAccel. Among the four childhood and adulthood circumstances domains, adulthood adversity was the largest contributor (9%), while adulthood SES, childhood adversity, and childhood SES accounted for 2.8%, 2.1%, 0.7%, respectively. All five domains of PGSs contributed 3.8% of variance in PhenoAgeAccel.

**Fig 1.**
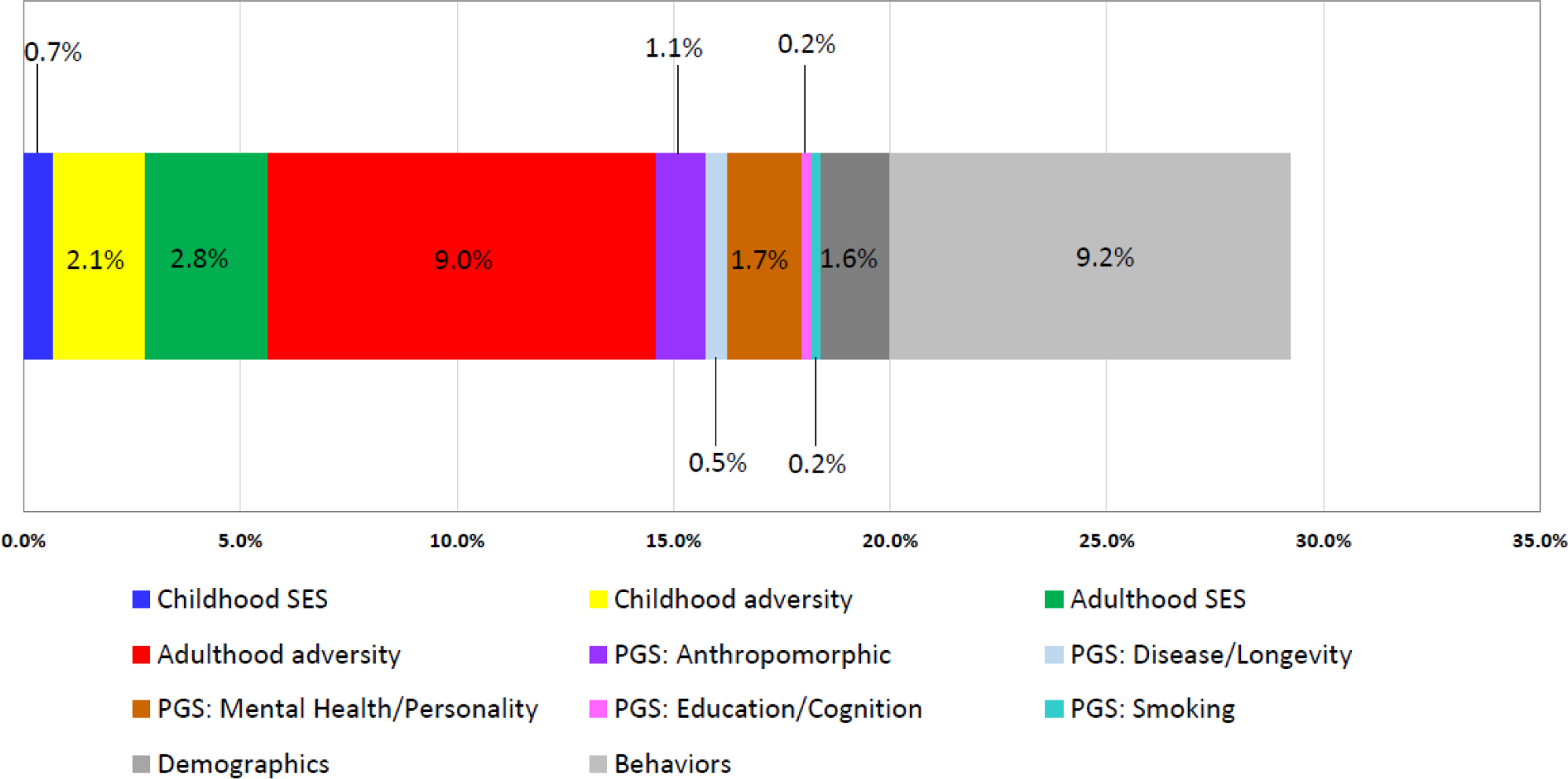
The contribution of all 11 domains to PhenoAgeAccel. SES, socioeconomic status; PGS, polygenic score. The 11 domains include four childhood and adulthood circumstances domains, five PGSs domains, demographics, and behaviors domains. Overall, the 11 domains contributed 29.2% (standard error = 0.003) of variance in PhenoAgeAccel.

### Profiles of childhood and adulthood circumstances and their relation to phenotypic aging

Using HCA, we identified six distinct subpopulations/clusters characterized by shared childhood and adulthood circumstances (Figure S2, top colored row). Figure 2A suggests that those assigned to the red cluster are characterized by having lower levels of education, having poor financial situations during childhood and adulthood, having lower educated parents, and living in neighborhoods with severe physical disorder. Conversely, the green cluster includes participants who had high adult SES, moderately high childhood SES, and who lived in neighborhoods with very low levels of physical disorder. The turquoise cluster includes participants with the highest SES in childhood and adulthood, but whose neighborhoods were slightly more disordered than those in the green cluster. Those in the yellow cluster are mainly characterized by having had higher levels of chronic stress, while those in the orange cluster had low SES in childhood and high levels of childhood trauma. Finally, the blue cluster represent participants with moderate SES in childhood, and very low levels of chronic stress. Figure S3 shows the correlation among the six clusters. Inverse relationship was observed between the green and red clusters—suggesting that participants assigned to them have opposite life experiences/circumstances. Similarly, the turquoise and orange appeared to represent opposite experiences, as did the yellow and the blue clusters.

Bivariate differences in PhenoAgeAccel between the six clusters are presented in Figure 2B. On average, the three clusters that appear to represent “disadvantaged” childhood and adulthood circumstances (i.e. red, yellow, and orange) exhibited higher phenotypic aging. Participants assigned to the red or yellow clusters were about 1.75 years older phenotypically than expected based on their chronological ages, while those in the orange cluster were about 0.3 years older than expected. Conversely, “advantaged” subpopulations/clusters, such as those in the green or the turquoise cluster had PhenoAge that were about two years younger than expected, while those in the blue cluster were a little over one year younger than expected.

**Fig 2.**
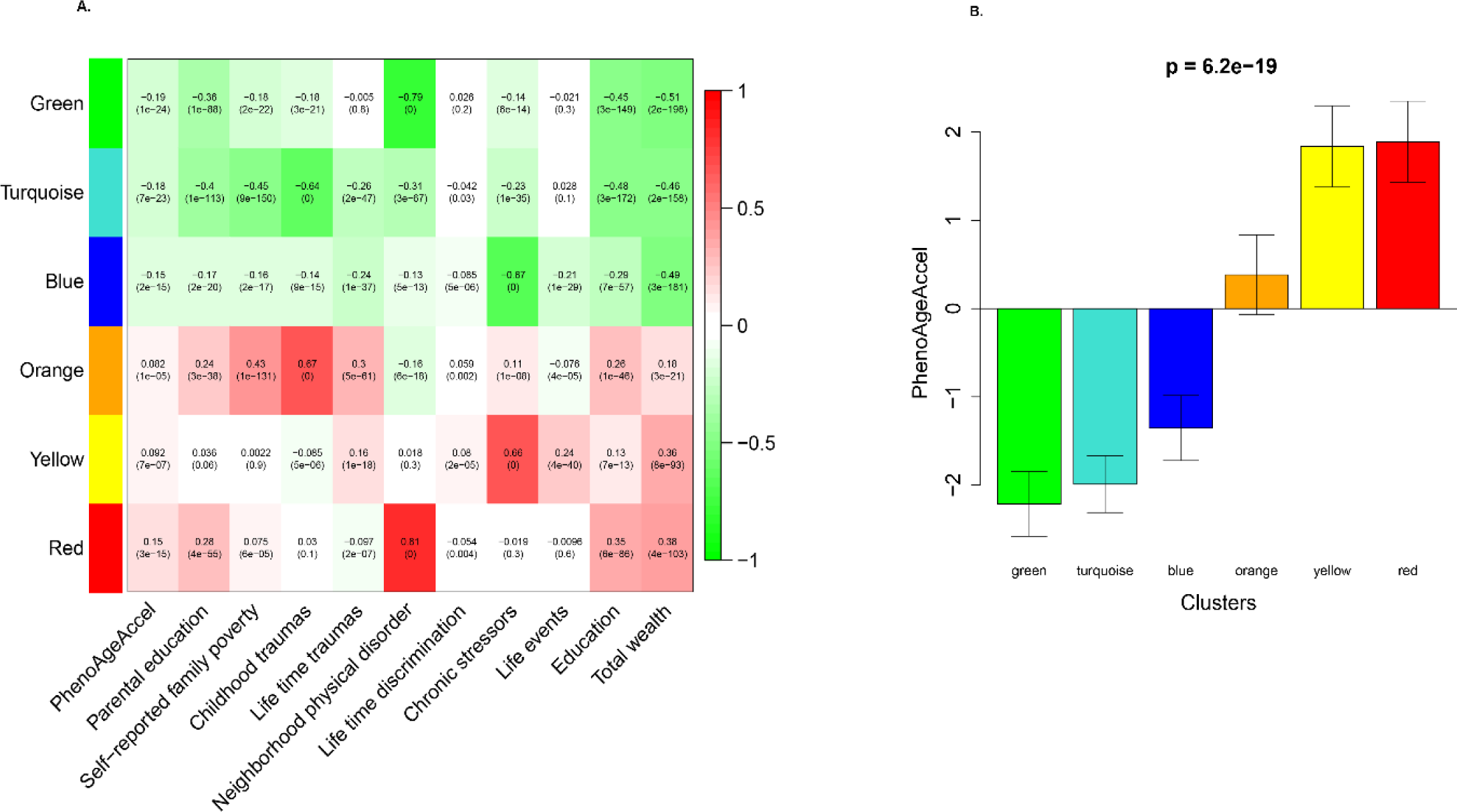
A) Cluster membership-traits correlations and p-values; B) PhenoAgeAccel across six clusters. In A), to determine what each subpopulation/cluster represents, we calculated a continuous measure (cluster membership) for each cluster (between −1 and 1) that denotes how strongly a person belongs to that given cluster—for instance, someone may have a score of 0.8 for the green cluster and −0.6 for the red cluster, suggesting he/she is very similar to the profile representative of the green cluster, but not the red cluster. Each cell reports the correlation (and p-value) resulting from correlating cluster membership (rows) to traits (columns, including PhenoAgeAccel and summarized measures of several circumstances). The table is color-coded by correlation according to the color legend.

### Gene-by-environmental interactions

Because PGS for coronary artery disease (CAD-PGS) accounted for the highest proportion of variance in PhenoAge (Figure S4), we tested the interaction between the six clusters and the CAD-PGS using OLS models. Significant main effects were found for both the circumstances clusters and PGS (Table 1). For instance, in a fully adjusted main-effect model, those assigned to the red cluster had PhenoAge that were more than 3.5 years older than those assigned to the green cluster (β=3.63, P=7.7E-9). Additionally, every one standard deviation increase in PGS was associated with a 0.44 year increase in PhenoAge (β=0.44, P=1.4E-2). In a subsequent interaction model, we observed a significant interaction between CAD-PGS and the red (relative to the green) cluster (β=1.50, P=1.9E-2). As illustrated in Figure 3, a higher CAD-PGS increased the difference in PhenoAge between those in the red versus the green cluster in a multiplicative manner, such that being in the red cluster versus the green cluster was associated with only a 0.6 year increase in PhenoAge for those with a PGS that was 2 standard deviations below the mean. However, among those with a PGS that was 2 standard deviations above the mean, being in the red cluster versus the green cluster was associated with a 6.6 years increase in PhenoAge.

**Table 1.** PhenoAge Associations with CAD-PGS and Childhood and Adulthood Circumstances.

|  | **Model 1** |  | **Model 2** |  |
| --- | --- | --- | --- | --- |
|  | Coef. (SE) | P Value | Coef. (SE) | P Value |
| **Six clusters for childhood and adulthood circumstances** |  |  |  |  |
| Green | Ref. |  | Ref. |  |
| Turquoise | 0.83 (0.58) | 0.151 | 0.84 (0.58) | 0.146 |
| Blue | -0.02 (0.61) | 0.967 | -0.01 (0.61) | 0.983 |
| Orange | 1.94 (0.66) | 0.003 | 1.96 (0.66) | 0.003 |
| Yellow | 3.46 (0.61) | 1.6E-8 | 3.50 (0.61) | 1.3E-8 |
| Red | 3.63 (0.63) | 7.7E-9 | 3.64 (0.63) | 7.0E-9 |
| **CAD-PGS** | 0.44 (0.18) | 0.014 | -0.09 (0.40) | 0.822 |
| **Gene-by-environmental interactions** |  |  |  |  |
| CAD-PGS*Green |  |  | Ref. |  |
| CAD-PGS*Turquoise |  |  | 0.44 (0.56) | 0.430 |
| CAD-PGS*Blue |  |  | 0.54 (0.58) | 0.354 |
| CAD-PGS*Orange |  |  | 0.66 (0.66) | 0.318 |
| CAD-PGS*Yellow |  |  | 0.41 (0.60) | 0.497 |
| CAD-PGS*Red |  |  | 1.49 (0.64) | 0.019 |

CAD-PGS, polygenic score for coronary artery disease. SE, standard error.
Model 1 adjusted for chronological age, sex, ancestry, proportion of experiencing obesity, and smoking. Model 2 additionally added the CAD-PGS\*clusters interaction term.

**Fig 3.**
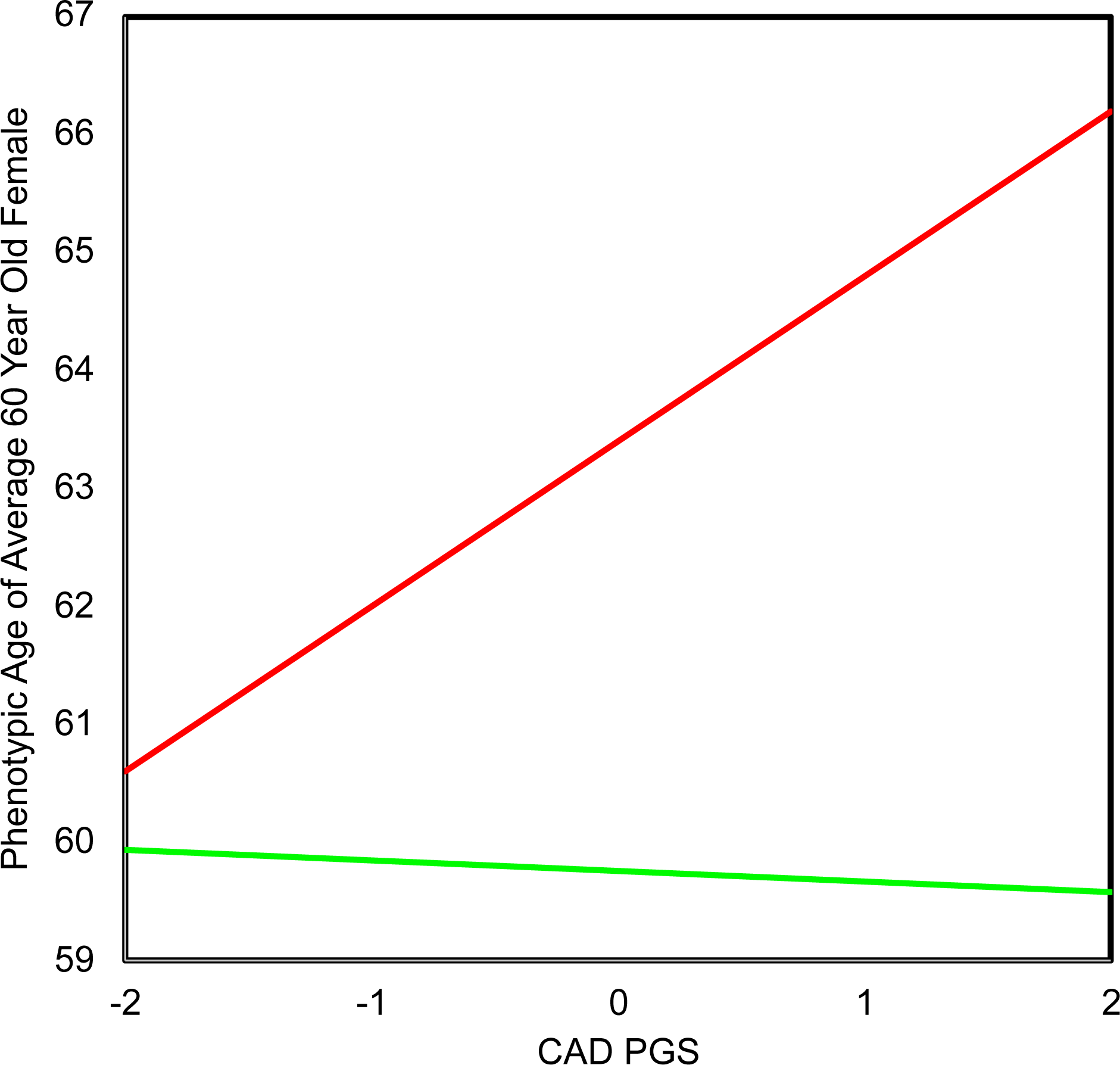
The significant interaction between CAD-PGS and the red (relative to the green) cluster for PhenoAge. CAD-PGS, polygenic score for coronary artery disease. This figure is based on an OLS model with adjustment for chronological age, sex, ancestry, obesity, and smoking, and the CAD-PGS*clusters interaction term.

## Discussion

Using data from a nationally representative study of older adults in the U.S., we showed that childhood and adulthood circumstances and genetic factors were associated with differences in a clinically-based signature of aging. The Shapley Value Decomposition approach revealed that the factors evaluated collectively accounted for under one-third of the variance in phenotypic aging. Furthermore, based on childhood and adulthood circumstances, we were able to group participants into subpopulations/clusters, which exhibited substantially different levels of phenotypic aging. Taken together, these results may inform potential interventions to reduce the health inequalities experienced throughout the life course. While causality needs to be formally evaluated, results from the current study highlight the socioenvironmental factors that potentially have the largest influence over levels of phenotypic aging. As such, targeting these factors may lead to improvements in health and diminish disparities.

Results from both the Shapley Value Decomposition approach and HCA draw attention to adversity in adulthood as a potential driver of differences in phenotypic aging and subsequent health disparities. More specifically, persons with higher levels of neighborhood physical disorder or high chronic stress—which were the most evident defining characteristics of the red and the yellow subpopulations/clusters—appear to phenotypically age faster than their peers. Our study extends results from earlier studies reporting that persons living in neighborhoods characterized by poor environments (e.g., lower aesthetic quality, and safety) exhibited accelerated cellular aging, proxied by shorter telomere length.^41 42^ Poor neighborhood environments may induce stress, predispose one to stressful life events, or shape exposure and vulnerability to stress.^43^ To date, both human and animal research have documented the negative effects of stress on health and aging.^44 45^ For instance, the literature on “allostatic load” postulates that stressful life experiences “get under the skin” and contribute to multi-system dysregulation. However, by considering multiple potential stressors simultaneously using advanced statistical approaches, we were able to assess their relative contributions. Our results suggest that preventing or reducing these adversities (e.g. through improvements in neighborhood safety and increasing affordability of housing) should be prioritized in efforts to improve population health, particularly in the face of rapid population aging in the U.S. and worldwide.

Similarly, our results add further evidence to substantiate the link between adult SES and aging. Education is thought to act as a robust indicator of SES, contributing to social gradients. In this study, the most defining shared attribute across the three advantaged subpopulation/clusters (i.e. green, turquoise, and blue) was higher levels of education. This is in stark contrast to the three disadvantaged clusters—all of which were characterized by low education. Chronic socioeconomic deprivation associated with low education is thought to provoke a number of adverse biological responses, including a gene expression profile called the conserved transcriptional response to adversity (CTRA), characterized by increased proinflammatory, signaling, and downregulation of antiviral type I interferon and antibody related genes.^46^ Over time, this transcriptional profile is thought to increase susceptibility to a number of age-related conditions, potentially via acceleration of the biological aging process.

As with adult SES, a large number of studies have highlighted the influence of childhood circumstances (e.g., SES and adversity).^10–17^ For instance, the “long arm of childhood” theory posits that childhood social and economic conditions get embedded within one’s biology and have far reaching implications for one’s health as s/he ages^47^ However, it is important to account for the fact that individuals who experience disadvantages in childhood are more likely to go on to experience adverse circumstances in adulthood, which could lead to overestimates of the effect of childhood circumstances in traditional regression analysis. In this study, we applied the Shapley Value Decomposition approach to appropriately decompose contributions of childhood and adulthood circumstances, providing relatively accurate estimates. We observed the influence of childhood SES and adversity on phenotypic aging, suggesting that adversity in childhood may influence aging beyond predisposing a person to adversity in adulthood.

While our results highlight the influence of SES and adversity on phenotypic aging, the detrimental effects were not consistent across individuals. Our results suggest a moderating effect of genetic predisposition, such that individuals who have an innate susceptibility to diseases, such as CAD, may suffer even more from experiences of chronic stress and adversity. We showed that for participants with low genetic risk for CAD, the differences in PhenoAge between those in the most adverse socioeconomic environments (red cluster) and those in the most advantageous (green cluster), were minimal—perhaps only ½ year. However, among individuals with high genetic risk, we observed a more than 6.5 year difference in PhenoAge between these two socioeconomic groups. This is noteworthy, given that we have previously shown that every 1-year increase in a person’s PhenoAge, relative to his/her chronological age, is associated with a 9% increased risk of dying.^8^ Based on these findings, it may be beneficial to target interventions and policies towards disproportionally at-risk individuals.

Despite the availability of a comprehensive set of circumstances variables and the innovative application of the Shapley Value Decomposition approach, the results should be interpreted with caution. First, the efforts to assemble a comprehensive set of circumstances from several sub-studies reduced sample size and potentially altered the population structure. For instance, a sizable proportion of persons who attended the 2016 VBS were excluded due to missing data on childhood adversity, collected through the 2015 LHMS. As a result, we observed slight differences in relevant characteristics of our analytic sample compared with those excluded in the analyses. This issue was partially offset by using dummy variables for missingness in order to retain participants, and/or considering other versions of survey weights in the sensitivity analyses—all of which produced findings consistent with those presented. Second, information on the specific timing of childhood and adulthood circumstances was not available. Previous studies have suggested that the earlier the adversity developed, the greater the negative effect on health in later life.^48^ Third, the five variables for major events in adulthood adversity domain were asked “before age 50”; therefore, we cannot rule out that they occurred in childhood. Fourth, most of these circumstances were based on self-reports, leading to possible recall biases, particularly as it relates to reporting of childhood experiences. In future research, it will be important to examine the associations between childhood adversity/SES and measures such as PhenoAge, to determine when individuals diverge in their aging rates. Similarly, longitudinal analyses that can test actual rate of change are also needed.

## Conclusions

In a national-representative sample of U.S. older adults, we demonstrated that socioenvironmental circumstances during both childhood and adulthood account for a sizable proportion of the difference in phenotypic aging. Furthermore, we provided evidence of a GxE interaction, suggesting that experiencing adverse circumstances may be more detrimental to individuals with a genetic predisposition to poorer health—in this case, increased risk of CAD. These findings may inform policy for allocating resources to promote healthy aging, and eventually ameliorate health disparities.

## Acknowledgements

This research was supported by NIH/NIA 4R00AG052604-02 (Levine), James Tobin Fund at Yale Economics Department (Chen), Yale Macmillan Center faculty research award (Chen), the PEPPER Center Scholar Award (P30AG021342) and other NIH/NIA grants (K01AG053408; R03AG048920) (Chen). Dr. Gill is the recipient of an Academic Leadership Award (K07AG043587) from the National Institute on Aging. No funder had any role in the study design; data collection, analysis, or interpretation; in the writing of the report; or in the decision to submit the article for publication.

